# *triangulaR:* an R package for identifying AIMs and building triangle plots using SNP data from hybrid zones

**DOI:** 10.1101/2024.03.28.587167

**Authors:** Ben J. Wiens, Jocelyn P. Colella

## Abstract

Hybridization provides a window into the speciation process and reshuffles parental alleles to produce novel recombinant genotypes. The presence or absence of specific hybrid classes across a hybrid zone can provide support for various modes of reproductive isolation. Early generation hybrid classes can be distinguished by their combination of hybrid index and interclass heterozygosity, which can be estimated with molecular data. Hybrid index and interclass heterozygosity are routinely calculated for studies of hybrid zones, but available resources for next-generation sequencing datasets are computationally demanding and tools for visualizing those metrics as a triangle plot are lacking. Here, we provide a resource for identifying ancestry-informative markers (AIMs) from SNP datasets, calculating hybrid index and interclass heterozygosity, and visualizing the relationship as a triangle plot. Our methods are implemented in the R package *triangulaR*. We validate our methods by simulating genetic data for a hybrid zone between parental groups at low, medium, and high levels of divergence. We find that accurate and precise estimates of hybrid index and interclass heterozygosity can be obtained with sample sizes as low as five individuals per parental group. We explore various allele frequency difference thresholds for AIM identification, and how this threshold influences the accuracy and precision of hybrid index and interclass heterozygosity estimates. We contextualize interpretation of triangle plots by describing the theoretical expectations for covariance of hybrid index and interclass heterozygosity under Hardy-Weinberg Equilibrium and provide recommendations for best practices for identifying AIMs and building triangle plots.

## Introduction

Research on hybridization provides insight into the evolution of reproductive isolation, the genetic basis of phenotypic variation, and novel modes of adaptation (Jones et al., 2018; Cronemberger et al., 2020; Aguillon et al., 2021; Nikolakis et al., 2022). Evolutionary outcomes of hybridization also often have implications for conservation and management of genetic diversity in natural populations (Chan et al., 2019). A common goal in studies of hybridization is to classify individuals as hybrids or as members of the parental groups (Gompert & Buerkle, 2013). Six hybrid classes are typically used to describe genetic variation in hybrid zones (Fitzpatrick, 2012): the first filial generation (F1), second filial generation (F2), backcrosses in each direction (BC), and the parental groups (P1, P2). Such classification serves as a starting point to describe hybrid zone dynamics and form hypotheses about the genetic, ecological, and environmental mechanisms that shape hybridization outcomes (Simon et al., 2018; Thompson et al., 2023).

Molecular data contain the necessary information for assigning hybrid classes (Lynch, 1991), which can be done by pairing hybrid index (i.e. ancestry proportions, or the proportion of alleles inherited from each parental group) with interclass heterozygosity (the proportion of loci with alleles from both parental groups). Visualizations of interclass heterozygosity (y axis) against hybrid index (x axis) are referred to as triangle plots, because the set of possible coordinates forms the shape of a triangle (Fig. 1). Hybrid index is measured in terms of proportion of ancestry from one parental group, such that individuals belonging to that parental group have a hybrid index of 1 and individuals of the other parental group have a hybrid index of 0. If only fixed differences between the parentals are used to calculate hybrid index, each parental individual, by definition, will have an interclass heterozygosity of 0. By the same logic, F1s will have an interclass heterozygosity of 1 and hybrid index of 0.5, assuming no gene conversion during recombination.

**Figure 1.**
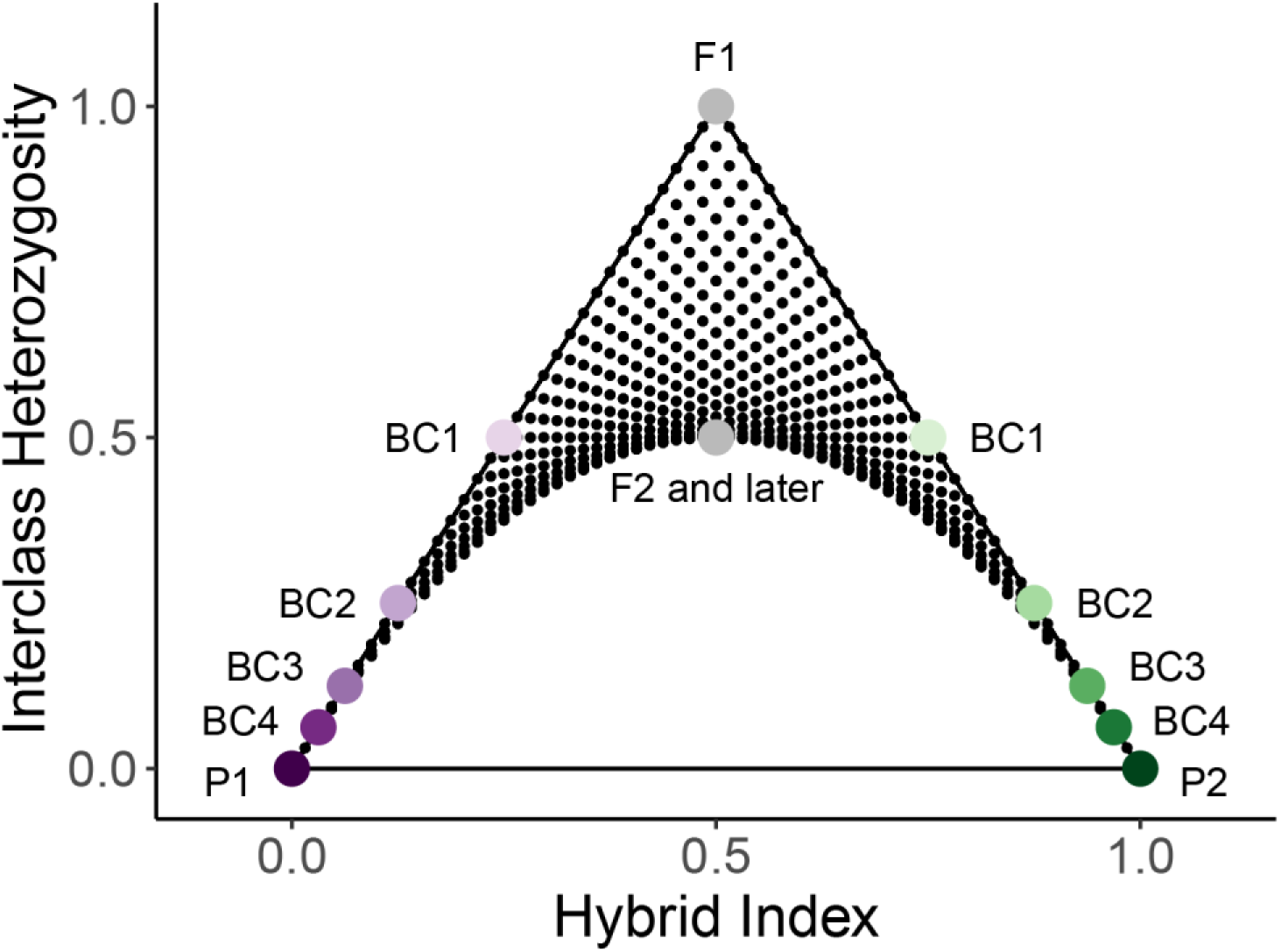
A triangle plot illustrating the theoretical expectations for combinations of hybrid index and interclass heterozygosity under Hardy-Weinberg Equilibrium (HWE). Large, colored points identify parental taxa (P1, P2) and hybrid classes (F1=first filial generation, F2=second filial generation, BC=backcross and the number of generations backcrossed toward the nearest parental population). Smaller black points show the possible combinations of hybrid index and interclass heterozygosity after six generations of mixing under HWE.

If including sites that are not fixed for alternate alleles in the parental groups, but which still show high differentiation, those values may not be matched exactly, but hybrid classes are still identifiable (Rosenberg et al., 2003; Fitzpatrick, 2012).

An early method (*introgress*) took a maximum likelihood approach to infer hybrid index and heterozygosity from codominant markers (e.g. amplified fragment length polymorphisms) or biallelic molecular markers (Buerkle, 2005; Gompert & Buerkle, 2009). More recently, Bayesian methods (*ENTROPY, bgc, bgchm*) have been developed to infer intersource ancestry and admixture proportions from SNP genotypes and genotype likelihoods (Gompert & Buerkle, 2012, 2013; Lindtke et al., 2012; Shastry et al., 2021; Gompert & Buerkle, *in review*). While those methods have provided substantial insight into the evolutionary ecology of hybridization, they also all impose a high degree of computational burden on the user, as they require unique file formats for the input data, implement time-intensive sampling of posterior distributions, and/or don’t provide intuitive visualizations of triangle plots. As such, there is a need for computational methods that are less intensive, which will facilitate quick descriptions of admixture and identification of hybrid classes when exploring new datasets.

When the genotypes for many (e.g. thousands) of biallelic sites are known with high confidence and parental populations are sampled, a simple analytical approach to ancestry estimation can be taken. With parental individuals included in the sample, ancestry-informative markers (AIMs) can be identified and used to calculate hybrid index and interclass heterozygosity. Whole genomes and reduced-representation sequencing methods, like RADseq and target capture, can provide many (e.g. thousands) SNPs from which AIMs can be identified, but practices for sampling parental populations and identifying AIMs vary widely (e.g. Del-Rio et al., 2022; Ocampo et al., 2023; Preckler-Quisquater et al., 2023). The sample size needed for each parental group is an important consideration because any method for identifying AIMs depends on identifying genomic sites that reliably show differentiation between the parental groups (Rosenberg et al., 2003). Short of sampling all individuals in a population, it is impossible to know the true allele frequencies, therefore, sample sizes must be large enough to provide reasonable estimates of allele frequencies in each parental population. Practically, the sample size must be large enough to produce AIMs that enable accurate and precise estimates of hybrid index and interclass heterozygosity, while balancing the expense (both in terms of time and money) of sampling and sequencing many individuals.

Another important consideration is the method used to identify AIMs. Intuitively, SNPs with fixed differences are most informative for estimating ancestry proportions, but for some genomic datasets restricting AIMs to such sites is not feasible (DeRaad et al., 2023). When there are not enough fixed differences between two parental groups, a lower threshold of differentiation must be used to identify AIMs. While FST is sometimes used as a metric of differentiation, it is susceptible to providing misleading results in cases when the two parental groups have different demographic histories, when there is low differentiation between parental groups, or when sample sizes are uneven, and it can be strongly influenced by within-group levels of variation (Meirmans & Hedrick, 2011; Jakobsson et al., 2013; Cruickshank & Hahn, 2014; Lotterhos & Whitlock, 2014;). Further, estimates of FST vary greatly when using different estimators (Berner, 2019). As an alternative to FST, the allele frequency difference (δ) between parental groups can be calculated for each site and used to set a threshold for defining AIMs (Rosenberg et al., 2003; Berner, 2019). Because restricting AIMs to sites with fixed differences is not realistic for all empirical datasets, it is worthwhile to explore the consequences of lower allele frequency difference thresholds for identifying AIMs, building triangle plots, and distinguishing between hybrid classes.

Here, we introduce the R package *triangulaR* (https://github.com/omys-omics/triangulaR), which provides simple and quick calculations of hybrid index and interclass heterozygosity from SNP datasets for building triangle plots and classifying hybrids. To validate these methods, we simulated genetic data for hybrid zones with parental groups at various levels of divergence. For each simulation, we investigate how the number of individuals sampled per parental group affects estimates of hybrid index and interclass heterozygosity. We find that sample sizes as low as five individuals per parental group provide reliable estimates of hybrid index and interclass heterozygosity, even when the parental groups are minimally divergent. We also show that when divergence is minimal, there is a practical tradeoff between restricting AIMs to only a few, highly informative sites versus relaxing the allele frequency difference threshold to include more, but less informative sites. We outline best practices for sampling parental populations, identifying AIMs, and building triangle plots to identify hybrids and hybrid classes from molecular datasets.

## Methods

### Package description

When parental populations are included in a sample from a hybrid zone and SNP genotypes are known with high confidence, hybrid index and interclass heterozygosity can be calculated analytically. Specifically, those metrics can be calculated by identifying AIMs, defined as sites with an allele frequency difference between the parental groups that is above a chosen threshold, and then polarizing alleles at those sites to determine in which parental group each allele has a higher frequency (Rosenberg et al., 2003).

This approach is implemented in the R package *triangulaR*. The main functions of this package are outlined in Table 1, and package documentation and a detailed tutorial are available at https://github.com/omys-omics/triangulaR. Briefly, *triangulaR* accepts as input genotype data for biallelic SNPs and leverages the functionality of the R package *vcfR* to read in vcf files and store vcfR objects (Knaus & Grünwald, 2017). The only other required input is an R dataframe with individual identifiers in the first column and population assignments in the second column. Hybrid index and interclass heterozygosity are calculated in two steps. First, SNPs with an allele frequency difference between the parental groups above a user-defined threshold are identified as AIMs, and a vcfR object containing individual genotypes at only these SNPs is returned. An allele frequency difference of 1 indicates a fixed difference between the two parental populations. In the second step, that vcfR object is used for calculating hybrid indices, which is done by summing the number of alleles in each individual that match one parental group and dividing by the total number of nonmissing sites present for that individual. Average interclass heterozygosity is also calculated during that step by counting the number of observed heterozygous sites and dividing by the total number of nonmissing sites. Dividing by the number of nonmissing sites is critical because dividing by all sites would essentially treat sites with missing data as homozygous and artificially deflate interclass heterozygosity estimates. Hybrid index and interclass heterozygosity estimates are returned along with user-defined population assignment and percent missing data for each individual, formatted in an R dataframe. Results can be visualized with a wrapper function (*triangle.plot*) that uses the R package *ggplot2* (Wickham, 2011) to plot the observed hybrid index and interclass heterozygosity values for each individual and draw the outline of the possible space on a triangle plot under HWE. Results can be colored by population or percent missing data to aid interpretation.

**Table 1.** Main functions of the R package *triangulaR*.

| Function | Description | Input | Output |
| --- | --- | --- | --- |
| `alleleFreqDiff()` | Identify AIMs from a SNP dataset. Implemented by filtering for SNPs that pass a given allele frequency difference threshold ($\delta$) | vcfR object;<br>popmap* | vcfR object<br>with AIM<br>genotypes |
| `hybridIndex()` | Calculate hybrid index and interclass heterozygosity for each individual | vcfR object<br>of AIMs;<br>popmap | R dataframe |
| `triangle.plot()` | Plot hybrid index and interclass heterozygosity of each individual on a triangle plot | R dataframe<br>(from<br>`hybridIndex()`<br>function) | Triangle plot |
| `missing.plot()` | Color individuals on triangle plot by percent missing data | R dataframe<br>(from<br>`hybridIndex()`<br>function) | Triangle plot |
| `AIMnames()` | Get the names of the SNPs that pass a given allele frequency difference threshold ($\delta$) | vcfR object;<br>popmap | vector of SNP<br>names |
| `specFreqDiff()` | Plot the distribution of allele frequency differences between the parental populations across all SNPs | vcfR object;<br>popmap | R dataframe |
| `aimFreqDist()` | Visualize the distribution of allele frequencies of AIMs in each parental population | vcfR object;<br>popmap | R dataframe |

\*R dataframe with individual IDs in the first column and population assignments in the second column

### Possible space on triangle plots under HWE

Triangle plots are used for identifying hybrid classes (e.g. F1s, backcrosses, etc.) and inferring the presence and strength of barriers to reproduction (Christe et al., 2016; Fitzpatrick, 2012; Lindtke et al., 2012; Pulido-Santacruz et al., 2018). It is therefore useful to consider the possible space on a triangle plot under Hardy-Weinberg Equilibrium (HWE). By this, we mean the possible combinations of hybrid index and interclass heterozygosity in the generations following interbreeding between two distinct parental groups, assuming random mating and the absence of selection, drift, or new mutations. Describing this space, therefore, provides neutral expectations for the possible combinations of hybrid index and interclass heterozygosity, against which observed combinations can be meaningfully compared.

When considering sites with fixed differences between parental groups, it follows that the hybrid index and interclass heterozygosity of an individual equate to the genome-wide frequency of the p1 allele and p12 genotype, respectively. Under HWE, genotype frequencies are given by allele frequencies, such that the frequency of the p12 genotype across an individual genome is calculated as 2(p1)(1 - p1), where p1 is the frequency of the p1 allele. Assuming HWE, if a cross occurs between two individuals with the same p1 allele frequency across the genome, the allele frequency will not change in the offspring and the genotype frequency of p12 is given by 2(p1)(1 - p1). Thus, on a triangle plot, the offspring will occur along the curve: p12 = 2(p1)(1 - p1). Alternatively, if a cross occurs between two individuals with different p1 allele frequencies, the expected genotype frequencies must be calculated by accounting for the variance in allele frequency between the two parents. Variance is calculated as:

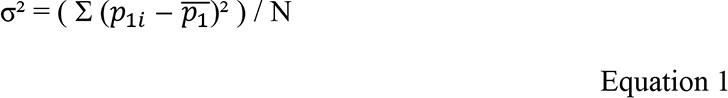

where p1*i* is the frequency of the p1 allele in the i^th^ parent, *p̅*_1_ is the average frequency of the p1 allele in the parents, and N is the number of parents (which is always two). The frequency of each offspring genotype is then:

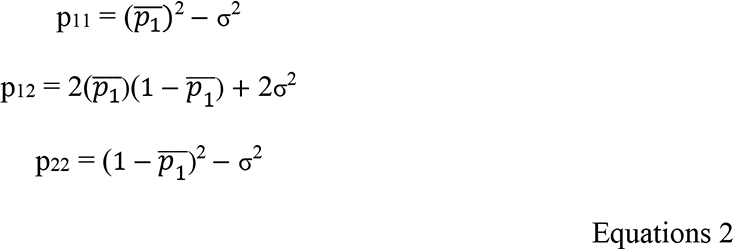

If both parents have the same p1 allele frequency, then the variance is 0 and the calculation of offspring genotype frequencies reduces to the standard formula under HWE. If the parents have different p1 allele frequencies, the hybrid index of the offspring will be the same as for a cross of two parents of the average p1 allele frequency, because the p11 and p22 genotypes will decrease by the same increment and changes in the frequency of the p12 genotype do not change the hybrid index. For offspring of parents with different p1 allele frequencies, the frequency of the p12 genotype will always be higher than for a cross of two parents of the average p1 allele frequency, because the variance is always positive. Therefore, under HWE, it is impossible for any cross to result in offspring below the curve defined by p12 = 2(p1)(1 - p1), because if the parents have the same allele frequencies their offspring will occur on that curve and if the parents have different allele frequencies their offspring will occur above it. To illustrate this point, we used Equations 2 to calculate and plot all possible combinations of hybrid index and interclass heterozygosity through six generations, starting with only the genotype frequencies of two parental individuals (Fig. S1).

### Simulations

To validate our methods for building triangle plots, we simulated genetic data for hybridization between two parental groups at three levels of divergence: low, medium, and high. We used SLiM 3 to perform forward-time genetic simulations under a non-Wright-Fisher model (Haller & Messer, 2019). Each simulation consisted of three phases (Fig. S2). Phase I lasted 1,000 generations and modeled a single, common ancestral population prior to divergence. Two populations existed during Phase I, with a high migration rate between them, such that in each generation 20% of individuals on average switched populations. Migration ceased during Phase II, which simulated allopatric divergence between the two parental populations. The length of Phase II differed for each simulation (low: 750, medium: 1000, high: 2000 generations), in order to reach different degrees of differentiation. Phase III modeled range expansion under a linear, stepping-stone model of 21 populations and lasted for 6,000 generations after contact was initiated between the two parental populations. The two parental populations were situated at the far ends of the simulation “landscape”, and migration could only occur between neighboring populations. Each generation, five migrants from each population with >200 individuals were randomly chosen to move into each adjacent population.

A 2 Mb diploid genome was simulated for each individual. Mutation rate was set to 10^-7^ per site per generation, all mutations were selectively neutral, and the recombination rate was 10^-5^ per site per generation. Those genomic parameters were chosen to generate SNP datasets on the order of magnitude (e.g. thousands) typically obtained from reduced-representation sequencing data, and such that linkage disequilibrium was minimal. Individuals were hermaphroditic, and could only reproduce if they existed in the previous generation. Each generation, every eligible individual reproduced with another randomly chosen individual in the same population. The number of offspring per mating followed a Poisson distribution (λ = 1.04). After all eligible individuals reproduced, they were removed from the simulation, and then migrants were randomly selected from among the offspring and moved to an adjacent population. Because individuals only existed for two generations and the carrying capacity per population was 1,100, the maximum number of eligible migrants per population fluctuated around 550.

At the end of Phase II, variant sites across all parental genomes were output in VCF format, and four hybrid classes (F1s, F2s, backcrosses in each direction) were simulated using custom R scripts. Specifically, 20 individuals from each parental population were randomly selected and paired with an individual from the other parental population, and 20 F1s were created by randomly choosing one allele from each parent at each genotype. Twenty F2s were created in the same way, but by pairing each F1 with another F1. Twenty backcrosses in each direction were created by pairing each F1 with a randomly chosen parental individual. From here on, we refer to this dataset as “known hybrids and parentals”.

During Phase III, random samples of 20 individuals from each population were taken every 200 generations for 6,000 total generations. Variant sites within the sampled individuals from each sampled generation were output in VCF format. The first sample was not taken until contact between the parental populations occurred. We define the first generation of contact as when all populations had more than 50 individuals. To ensure that contact happened in the central-most population (p10), migration could only occur into p10 once both adjacent populations (p9 and p11) contained more than 200 individuals. In this way, the first sample always occurred within the first few generations of hybridization in p10. We refer to that sample as generation 0, and subsequent samples as the number of generations since the first sample. At generation 0, we recorded the sites that contained fixed differences between all individuals of the parental populations, as a way of tracking the true degree of introgression over time.

### Number of individuals sampled from parental populations

The accuracy of observed allele frequencies in each parental population depends on the number of individuals sampled. When an allele occurs at low frequency in a population, the probability of detection increases as the sample size increases. We tested how sample sizes influence the number of AIMs that appear as fixed differences. Using the simulated dataset of known hybrids and parentals, we calculated the difference in allele frequency between parental populations at every variable site. Because all parental individuals were sampled for this dataset, these are the true allele frequency differences between the parental populations. We then randomly downsampled 20, 10, 5, and 2 individuals from each parental population, and used only those samples of individuals to calculate allele frequency differences. We repeated this procedure 200 times for each sample size. We report the distribution of true allele frequency differences at sites that appear to have fixed differences based on each replicated sample size. We then randomly chose one replicate of each sample size with which to calculate hybrid index and interclass heterozygosity of the hybrids and sampled parentals, based on sites with apparent fixed differences (δ=1).

### Allele frequency difference thresholds

For the dataset of known hybrids and parentals, we identified AIMs and built triangle plots. We tested three allele frequency difference thresholds for identifying AIMs: δ=1 (i.e. fixed differences), δ=0.75, and δ=0.5. Because this dataset includes all parental individuals in the simulation at the sampled generation, observed allele frequencies in each parental population are the true allele frequencies. Using AIMs that passed each allele frequency difference threshold, we calculated hybrid index and interclass heterozygosity for every hybrid and parental individual and built triangle plots.

### Accuracy and precision of estimates

To quantify the correctness of hybrid index and interclass heterozygosity estimates produced with different sample sizes from parental populations and different allele frequency difference thresholds, we calculated accuracy and precision for four hybrid classes (F1, F2, and both backcrosses), separately. Accuracy is defined as the difference between the expected value and the average observed value, divided by the expected value. We subtract this value from 1 to report percent accuracy.

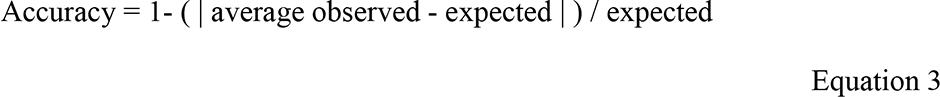

Precision is defined as the average absolute Euclidean distance of each individual estimate from the average estimate.

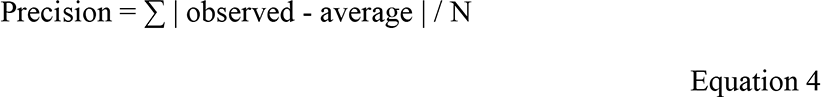

Precision is not subtracted from 1 or divided by the expected value, therefore the units reflect the Euclidean distance on the triangle plot, with smaller values indicating higher precision.

### Introgression and identifying AIMs

Identifying AIMs relies on the presence of diagnostic sites across the genomes of the parental groups. To investigate how introgression between parental groups influences AIM identification and calculations of hybrid index, we analyzed generations 0 through 6,000 of the simulated data. To track introgression over time, we identified sites with fixed differences in the parental populations (p0 and p20) at generation 0. We refer to these sites as “true” AIMs because they represent real differences prior to gene flow. We then created two additional sets of AIMs for each sampled generation by identifying sites with fixed differences (δ=1) and sites with allele frequency differences above 0.75 between the samples of the parental populations. In this way, we tested the accuracy of inferred hybrid indices in the face of introgression into the parental populations over time. For each set of AIMs, we calculated the average hybrid index of each population for generations 0 through 6,000.

## Results

### Simulations

At the end of Phase II (allopatric divergence), the low, medium, and high divergence simulations contained 6251, 6511, and 6894 variable sites, respectively. The parental populations were sampled at this point to create the dataset of known hybrids and parentals. There were 15 (low), 36 (medium), and 297 (high) fixed differences between the parental populations (Fig. S3). The average allele frequency difference between the parental populations following allopatric divergence was 0.10 (low), 0.11 (medium), and 0.16 (high). We calculated dXY and π in *pixy*, including invariant sites during calculation (Korunes & Samuk, 2021). Values of dXY between the parental populations were 0.00032 (low), 0.00036 (medium), and 0.00055 (high). Nucleotide diversity within each parental population was consistent across simulations (π=0.0002 ± 8.0x10^-6^). When contact began during Phase III, there were 28 (low), 59 (medium), and 345 (high) fixed differences between the parental populations. Although the average allele frequency difference and dXY between parental populations were not substantially different between the low, medium, and high divergence simulations, these levels of divergence provided very few to many fixed differences, which was our main consideration for evaluating the effects of sample size and allele frequency difference threshold on AIM identification.

### Number of individuals sampled from parental populations

As expected, the accuracy of estimated allele frequency differences increased as the sample size of the parental populations increased (Table S1). With a sample of 20 individuals per parental population the average accuracy of estimated allele frequency differences was >99% for all levels of divergence. A sample size of five per parental population also yielded high accuracy, at 93%, 95%, and 98% for low, medium, and high levels of divergence, respectively. The distribution of true allele frequency differences at sites with fixed differences in the sample are consistently left-skewed, with a peak at one (Fig. 2). With a sample size of five per parental population, 95% of sites with δ = 1 in the sample have a true allele frequency difference of at least 0.72 for all levels of divergence.

**Figure 2.**
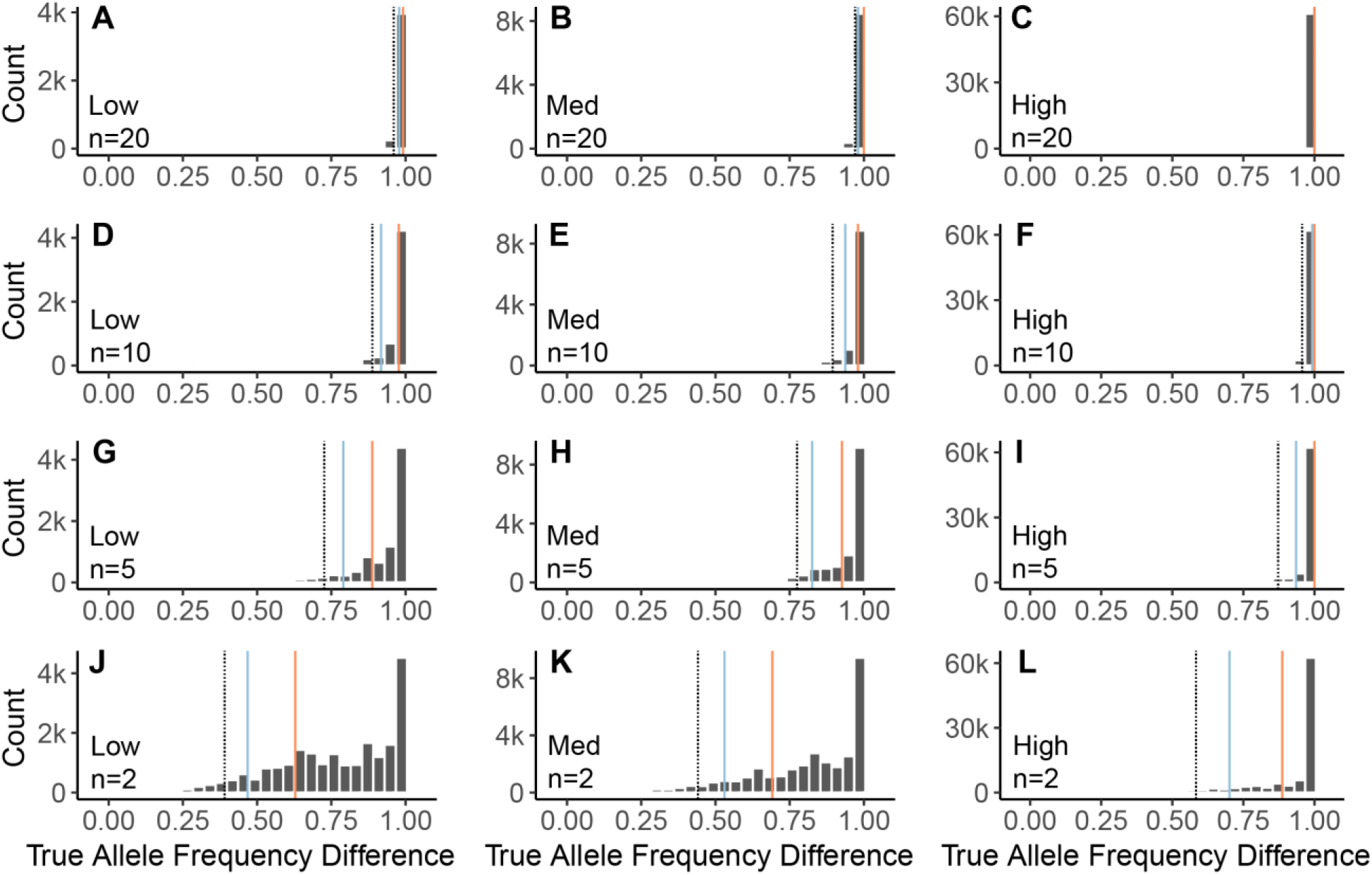
Distributions of the true allele frequency differences of sites that appear to have fixed differences with smaller parental sample sizes: 20 **(A-C)**, 10 **(D-F)**, 5 **(G-I)**, and 2 **(J-L)**. The observed distributions were created with 200 random sampling replicates of 20, 10, 5, or 2 individuals (n) from each parental population. The left column shows the simulation with low differentiation, center shows the simulation with medium differentiation, and the right column shows the simulation with high differentiation. On each plot, 95% of observed values fall above the black dotted line, 90% of the values fall above the blue solid line, and 75% fall above the red solid line.

The number of AIMs (δ = 1) identified increased as sample sizes decreased, due to an increase in false positive fixed differences; that is, differences that appear fixed in the sample of the parental groups but are not truly fixed in the populations (Fig. S4). Yet, even with a sample size of five from each parental population, hybrids appear as expected on triangle plots regardless of the level of divergence between parental populations. Estimated values of hybrid index and interclass heterozygosity for each hybrid class remained highly accurate (>90%) down to a sample size of five individuals from each parental population (Fig. S5). In general, accuracy increased as the sample size and the level of divergence increased. Estimated values of hybrid index and interclass heterozygosity were also highly precise (<0.1 average deviation) for all sample sizes and levels of divergence. Perhaps counterintuitively, for all hybrid classes except F1s, estimates became more precise as parental sample sizes decreased.

### Allele frequency difference thresholds

The number of AIMs identified increased as the allele frequency difference threshold decreased (Fig. 3). Estimated values of hybrid index and interclass heterozygosity for hybrids were generally most accurate when δ = 1, except for F2 estimates, which were more accurate when δ=0.75 (Fig. 4). Estimated values were generally very precise (<0.07 average deviation), with the exception of interclass heterozygosity estimates for F2s and backcrosses when using δ=1. For all hybrid classes except F1s, estimates became more precise as δ decreased. For F1s, estimates were most precise for δ = 1. In the case of δ = 1, parentals and F1s all had the exact expected values of hybrid index and interclass heterozygosity, regardless of the level of divergence.

**Figure 3.**
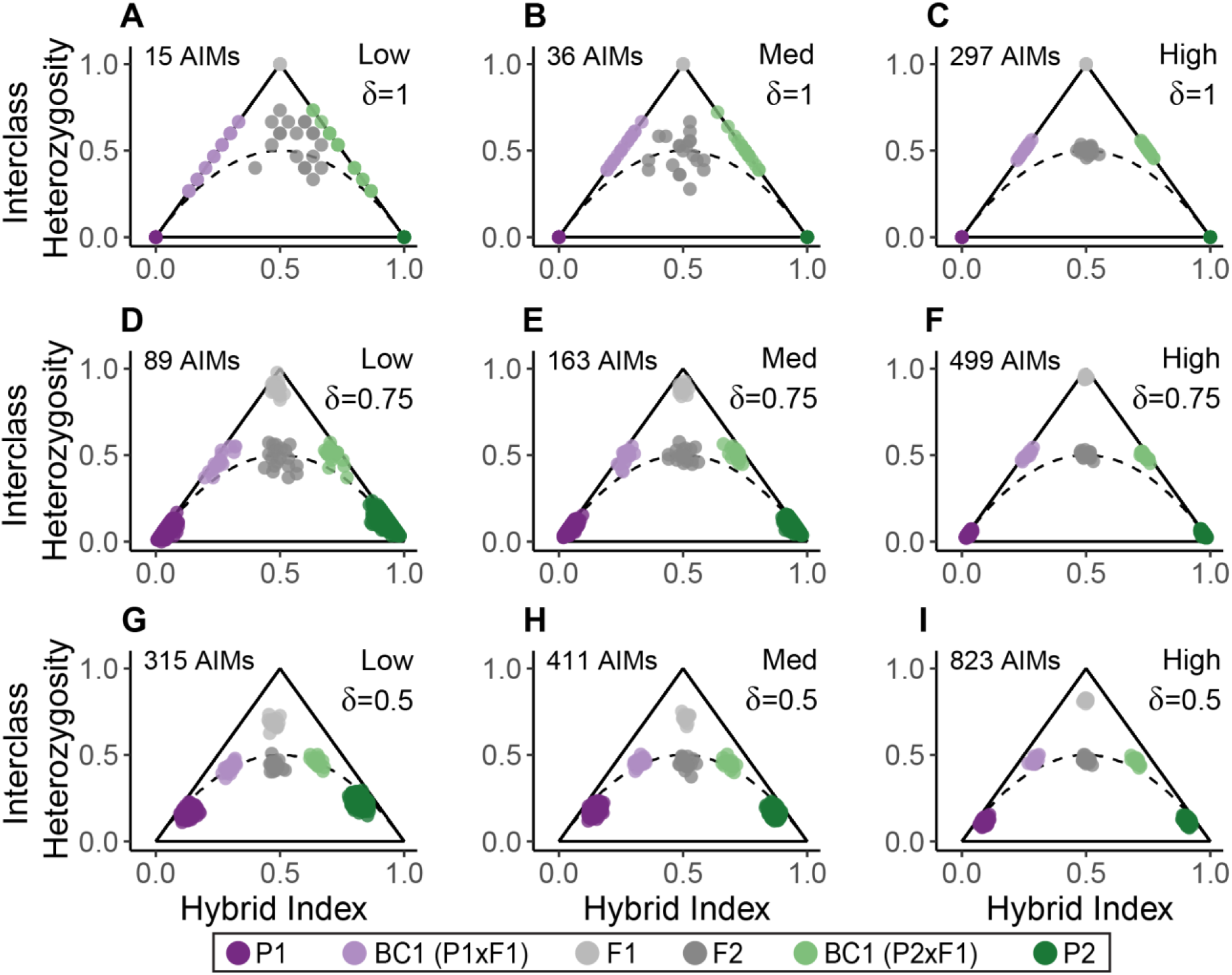
Triangle plots for known hybrids and parentals based on AIMs identified with true allele frequency difference thresholds (δ) of 1 **(A-C)**, 0.75 **(D-F)**, and 0.5 **(G-I)**. The left column shows the simulation with low differentiation, the center column shows the simulation with medium differentiation, and the right column shows the simulation with high differentiation. Solid black lines indicate the possible space on a triangle plot, and the dotted black curve indicates the boundary below which individuals cannot occur, assuming Hardy-Weinberg Equilibrium.

**Figure 4.**
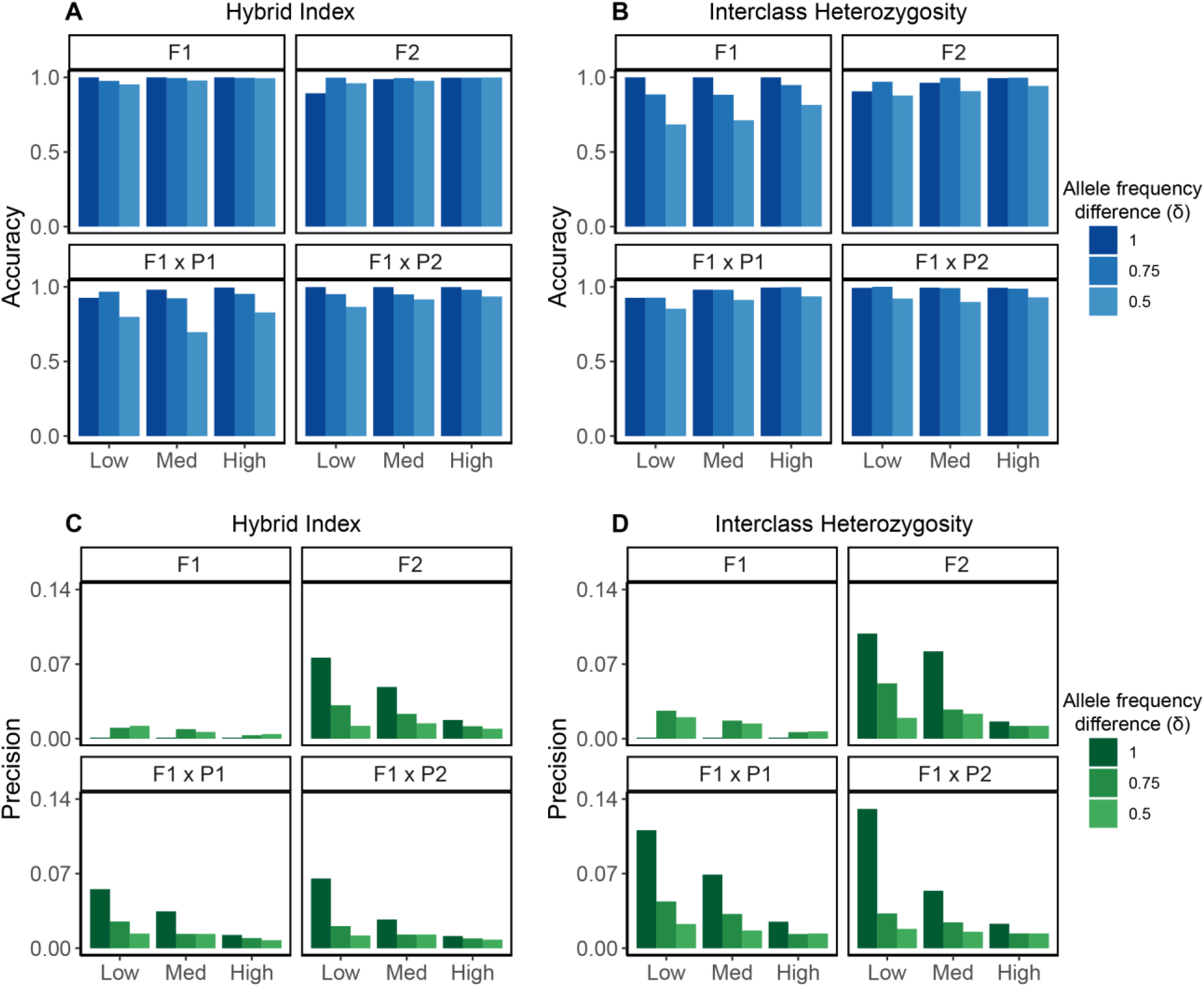
Accuracy **(A&B)** and precision **(C&D)** of hybrid index **(A&C)** and interclass heterozygosity **(B&D)** estimates based on AIMs identified with true allele frequency difference thresholds of 1, 0.75, and 0.5. Each simulation (low, medium, high) is shown on the x-axis. Accuracy and precision was measured for 20 individuals from each of the four hybrid classes (F1, F2, and the two first generation backcrosses) separately. Accuracy is reported as a percent, with 1 indicating 100% accuracy. Precision is reported as the average Euclidean distance of each observation within a class from the average of that class, such that smaller values indicate higher precision.

### Introgression and identifying AIMs

In each simulation, some alleles at sites that had fixed differences at the beginning of contact had introgressed into both parental populations (p0 and p20) by generation 1,000 (Fig. S6). By generation 6,000, each parental population contained at least 25% ancestry from the other parental population at sites that had started as fixed differences. Yet, the estimated ancestry of each parental population using AIMs (δ = 1) identified with 20 sampled individuals remained at 0 and 1 for p0 and p20, respectively, across all generations. Lowering δ to 0.75 resulted in the recognition of some admixture in the parental populations, but did not recover true levels of introgression into the parental populations.

## Discussion

Triangle plots provide a simple but powerful way to visualize genetic variation across hybrid zones and identify hybrids. Current approaches for building triangle plots are tailored for investigation of the evolutionary processes driving different patterns of ancestry across the genome (Gompert & Buerkle, 2012; Shastry et al., 2021). While these approaches are needed, they are computationally demanding, creating a need for simpler approaches designed for intuitive descriptions of hybrid zones. Further, a set of best practices for selecting sample sizes from parental populations and setting the allele frequency difference threshold for AIM identification are needed. Here, we present *triangulaR* (https://github.com/omys-omics/triangulaR), an R package for identifying AIMs, calculating hybrid indices and interclass heterozygosity, and visualizing triangle plots from SNP data. To facilitate use of *triangulaR* and interpretation of results, we simulated data to examine how two common criteria for identifying AIMs, the sample size of parental groups and the allele frequency difference threshold, influence estimation of hybrid index and interclass heterozygosity. Further, we provide a set of expectations for the covariance of hybrid index and interclass heterozygosity under HWE, against which empirical data can be compared. We then outline how observed combinations of hybrid index and interclass heterozygosity can be interpreted as support for common hybrid zone models. We anticipate *triangulaR* and the theoretical framework developed here will be useful for identifying hybrids, assigning individuals to hybrid classes, designing sampling schemes for natural populations, analyzing next-generation sequencing data, and interpreting triangle plots.

### How to choose an allele frequency difference threshold for AIM identification

There is a tradeoff between setting a high allele frequency difference threshold, which will recover fewer, more informative sites, versus selecting a lower threshold that will recover more, potentially less informative sites. The optimal threshold will depend on the study system, but as a best practice various thresholds should be explored during the analysis of empirical data. Our simulations suggest that the most important factor for accuracy and precision of hybrid index and interclass heterozygosity estimates is the number of AIMs used for calculations (Figs. 4 & S5). This is especially true when the level of divergence between parental groups is low. For example, using only the 15 fixed differences present between the parental populations in the low divergence simulation, we obtain perfect estimates of hybrid index and interclass heterozygosity for parentals and F1s, but imprecise estimates for F2s and backcrosses. When the allele frequency difference threshold is relaxed to 0.75, almost six times as many AIMs (N = 89) are recovered, and the precision and accuracy of hybrid index and interclass heterozygosity estimates increase for F2s and backcrosses, with minimal decrease in accuracy or precision of the estimates for F1s. F2 and backcross estimates are more sensitive to the number of AIMs used because there are three (p11, p12, p22) and two (p11, p12) possible genotypes at AIMs in these genomes, respectively, while there is only one genotype (p12) expected at AIMs in F1 genomes.

Additional patterns emerge when comparing triangle plots built for the same individuals but with AIMs called at different allele frequency difference thresholds (Fig. 3). For F1s, the accuracy and precision of interclass heterozygosity estimates decrease as the threshold decreases, accompanied by a downward shift in the estimated values themselves, while the accuracy and precision of hybrid index estimates are largely unaffected. For backcrosses, interclass heterozygosity estimates remain accurate and become more precise as the threshold decreases, but estimated values of hybrid index tend to shift towards the center of the plot (0.5). Taken together, a trend emerges wherein all hybrid classes gravitate towards the center of the triangle plot at lower allele frequency thresholds. This pattern becomes less pronounced when divergence between parental groups is higher, but should be taken into consideration when qualitatively inferring hybrid classes based on a triangle plot.

When analyzing empirical data, it is best practice to try multiple allele frequency difference thresholds and compare the resulting triangle plots. If many (>100) AIMs are recovered with δ = 1 and the resulting triangle plots are mostly unchanged with lower values of δ, then choosing δ = 1 is likely appropriate. Even as few as 30 fixed differences can produce reliable triangle plots, but selecting a lower threshold (e.g. δ = 0.75) may produce more precise estimates without sacrificing accuracy. When there are very few (<15) to no fixed differences between the parental populations, lower thresholds that provide more AIMs will be necessary to distinguish hybrid classes. We emphasize, however, that the reason lower thresholds can provide better estimates is because they increase the total number of AIMs. If few (<50) AIMs are identified even with a low allele frequency difference threshold (e.g. δ = 0.5), accurate and precise hybrid index and interclass heterozygosity estimates will be difficult or impossible to obtain (Figs. S7 & S8).

### What sample size of parental groups is needed?

To identify AIMs with which to calculate hybrid index and interclass heterozygosity, samples from both parental groups are required. The goal is to sample individuals that have not experienced admixture, such that they accurately reflect allele frequencies in the respective parental populations. Thus, an important practical consideration in experimental design is the number of parental individuals needed to provide reasonably accurate estimation of allele frequency differences between populations.

We describe the distribution of true population-wide allele frequency differences of AIMs that falsely appear as fixed differences in the parental group samples. When divergence between the parental groups is low, it is difficult to minimize the proportion of false positive fixed differences. In our simulation with the least divergence between parental populations (true fixed differences at only 0.2% of variable sites), a sample size of 20 individuals per parental population and δ = 1 resulted in a 29% false positive rate. That means that of the sites that appeared to be fixed for different alleles in the sample, 29% were not actually fixed in the population. Yet, it is important to consider the distribution of true allele frequencies of those perceived fixed differences. Sites with a high allele frequency difference between the parentals still provide valuable information, even if they are not fixed. With a sample size of 20 individuals from each parental population, >95% of identified AIMs (δ = 1) had a true allele frequency difference >0.95 for all simulated levels of divergence (Fig. 2, Table S1). Across all sample sizes, the distribution of true allele frequency differences at sites that appear to be fixed differences based on the sample is always left-skewed, meaning that the noise caused by the inclusion of some sites with a low allele frequency difference is minimal.

Counterintuitively, our simulations suggest that a sample size of five from each parental population can provide more precise estimates of hybrid index and interclass heterozygosity than larger sample sizes, without sacrificing accuracy (Fig. S5). That is because more sites pass the δ = 1 threshold when the parental sample size is smaller, resulting in a larger set of AIMs. While some of those sites are false positives in the sense that they are not truly fixed differences, they still exhibit large allele frequency differences between the parental populations (Fig. 2). As such, the precision is increased not because of the lower sample size *per se*, but through the inclusion of additional informative sites obtained with the lower allele frequency difference threshold. This is related to the consideration of δ as discussed above, whereby more precise and equally accurate estimates of hybrid index and interclass heterozygosity can be obtained by relaxing δ to increase the number of AIMs recovered.

In empirical systems, the optimal number of parental individuals to sample will depend on the question, level of divergence, and sequencing effort, among other variables. For deeply diverged species, hybrid index and interclass heterozygosity can be accurately estimated with as few as two or three samples from each parental species, whereas more shallowly diverged groups may require a parental sample size of 20 or more. But as a general rule of thumb, a sample of five individuals from each parental population will usually be sufficient to address questions related to hybridization, provided there is enough sequencing effort to obtain thousands of SNPs. In our simulations, the average accuracy of estimated allele frequency differences was >93%, even in the simulation with the least divergence between the parental populations. Further, even when divergence between parentals was low, sampling five individuals from each population resulted in 95% of identified AIMs (δ = 1) having a true allele frequency difference of at least 0.72 (Table S1). We also note that in our simulations, decreasing the parental sample size from twenty to five only marginally affected the accuracy of hybrid index and interclass heterozygosity estimates and precision actually increased for F2s and backcrosses. Hybrid classes also appeared as expected on triangle plots with parental sample sizes as few as five individuals (Fig. S4), an important consideration, as hybrid classes are often inferred qualitatively based on the combination of hybrid index and interclass heterozygosity (Fitzpatrick, 2012).

### Interpreting triangle plots

Triangle plots are not only useful for identifying hybrids and assigning hybrid classes, but have also been used as support for the presence/absence of barriers to reproduction (Fitzpatrick, 2012; Lindtke et al., 2012; Christe et al., 2016; Pulido-Santacruz et al., 2018). To help guide such inferences, we outline a theoretical framework for triangle plot interpretation based on the expectations of the covariance of hybrid index and interclass heterozygosity under HWE. We demonstrate mathematically that under HWE, the possible space on a triangle plot is defined by the lines y = 2x and y = -2x + 2, and the curve y = 2x(1 - x) (Fig. 1). Deviation from those expectations – most notably, individuals falling below the curve – is consistent with violation of HWE and can therefore provide support for the presence of barriers to reproduction, nonrandom mating, natural selection, and/or genetic drift (Pulido-Santacruz et al., 2018). For example, drift in an admixed population could cause individuals to occur below the curve defined by HWE through the random fixation of alleles over time. If alleles from either parental group have an equal chance of reaching fixation in the hybrid population, then interclass heterozygosity would decrease, but hybrid index would remain unchanged. Alternatively, support for post-zygotic isolation due to Dobzhansky-Muller incompatibilities (DMIs) may be gained due to the absence of F2s (or any individuals with hybrid index and interclass heterozygosity of 0.5), because DMIs are expected to manifest in the F2 generation, when there is no longer at least one copy of each parental allele at every locus (Maheshwari & Barbash, 2011; Thompson et al., 2023). Importantly, the absence of individuals below the curve defined by HWE is not evidence for barriers to reproduction or natural selection for or against hybrids; on the contrary, such an absence is expected when there is random mating and no selection.

Consideration of the expectations for triangle plots under HWE also informs inference of hybrid classes. F1s and first generation backcrosses are easily distinguished because their combination of hybrid index and interclass heterozygosity is unique. F2s, however, are indistinguishable from F3s and further crosses, because under HWE all are expected to have a hybrid index and interclass heterozygosity of 0.5. We also show that after only four generations of unidirectional backcrosses to one parental group, hybrid index (0.03125) and interclass heterozygosity (0.0625) will be only marginally different from members of the backcrossing parental group, making distinctions based on these metrics difficult.

### Caveats and other considerations

Our method performs well even when there is low divergence between parental groups, provided that there is enough sequencing effort across the genome to identify >30 SNPs with high allele frequency differences. We expect that in most cases, modern sequencing methods (e.g. RADseq, target capture) will provide large enough SNP datasets for this endeavor. However, if divergence between parental groups is extremely low, such that there are no fixed differences and <50 SNPs pass an allele frequency different threshold of 0.5, calculating hybrid index and interclass heterozygosity becomes inaccurate and imprecise. To overcome such limitations, a larger proportion of the genome will need to be sequenced. Sequencing more parental individuals and/or sequencing the same loci at higher coverage will filter out less-informative AIMs, but will not increase the number of AIMs identified, which is the primary driver of accuracy and precision of hybrid index and interclass heterozygosity estimates.

When interpreting triangle plots, it is important to consider the assumptions made when choosing an allele frequency difference threshold and assigning individuals to parental groups. When using only fixed differences (δ = 1) as AIMs, all individuals assigned as parentals will, by definition, have an interclass heterozygosity of 0 and hybrid index of either 0 or 1. An individual misassigned to either parental group will, therefore, be impossible to diagnose in this context. Thus, even if there are enough fixed differences to justify a threshold of δ = 1, it is still worthwhile to explore lower thresholds, with which it may be possible to distinguish misassigned parentals from true parentals. That is because the interclass heterozygosity and hybrid index of individuals assigned to the parental groups can vary at lower allele frequency difference thresholds. Thus, misassigned parentals will exhibit more intermediate hybrid index and higher interclass heterozygosity than true parentals. To corroborate hybrid indices calculated with lower allele frequency difference thresholds, ancestry proportions can be inferred with unsupervised clustering algorithms, such as those implemented by *STRUCTURE* (Pritchard et al., 2000) or *ADMIXTURE* (Alexander et al., 2009), although care should be taken to avoid common misinterpretations of results from these approaches (Bradburd et al., 2018; Lawson et al., 2018).

More problematic is if there are no true parentals in the sample. In the absence of diagnostic morphological features, it may not be possible to detect introgression into parental populations. In our simulations, introgression occurs into the parental populations by generation 1,000, but it is impossible to detect using allele frequency data from that generation on, because sites that began as fixed differences between the two parental groups now occur in both groups. As such, sites that were once informative of ancestry become indistinguishable from shared ancestral variation (Fig. S6). The true history of introgression is therefore obscured by the unknowability of ancestral allele frequencies. Other work has shown that admixture proportions inferred by unsupervised cluster algorithms also face this problem, because the models cannot distinguish admixture in every individual from shared ancestral variation, and subsequently interpret the least-admixed individual(s) to belong entirely to one genetic cluster (Lawson et al., 2018).

### Future directions

There remains a need for methods that can accurately estimate admixture proportions and interclass heterozygosity in cases where parental groups are minimally divergent or have experienced introgression. In the case of introgression into parental populations, analyzing the distribution of coalescent heights across the genome may prove effective for distinguishing introgressed loci from shared ancestral variation (Hibbins & Hahn, 2022). Additional work is also needed to develop the theoretical expectations for observed combinations of hybrid index and interclass heterozygosity in the presence of pre- and/or post-zygotic reproductive isolation. We expect that *triangulaR* will be a useful community resource for identifying and describing hybridization using genomic data, and that with continued theoretical development, triangle plots will serve as an effective tool for understanding the evolutionary processes governing hybrid zone dynamics.

## Acknowledgements

We thank Devon DeRaad and Lucas DeCicco for insightful discussions and Marlon Cobos for guidance in R package development. This work was supported by the HPC facilities operated by the Center for Research Computing at the University of Kansas. This work was supported by a National Science Foundation award to PI Colella (NSF#2100955).

## Author contribution statement

BJW conceptualized the study, wrote the R package *triangulaR*, performed the analyses, generated the figures, and drafted the manuscript. JPC funded the project, provided conceptual feedback, and contributed to the editorial process.

## Data archiving

The R package *triangulaR* is available from https://github.com/omys-omics/triangulaR. SLiM, Bash, Python, and R scripts used to simulate genetic data and perform analyses are available on GitHub at https://github.com/omys-omics/triangle_plot_sims. Raw VCF output from SLiM simulations are available at the same link.

