## Supplemental Information for "*triangulaR:* an R package for identifying AIMs and building triangle plots using SNP data from hybrid zones"

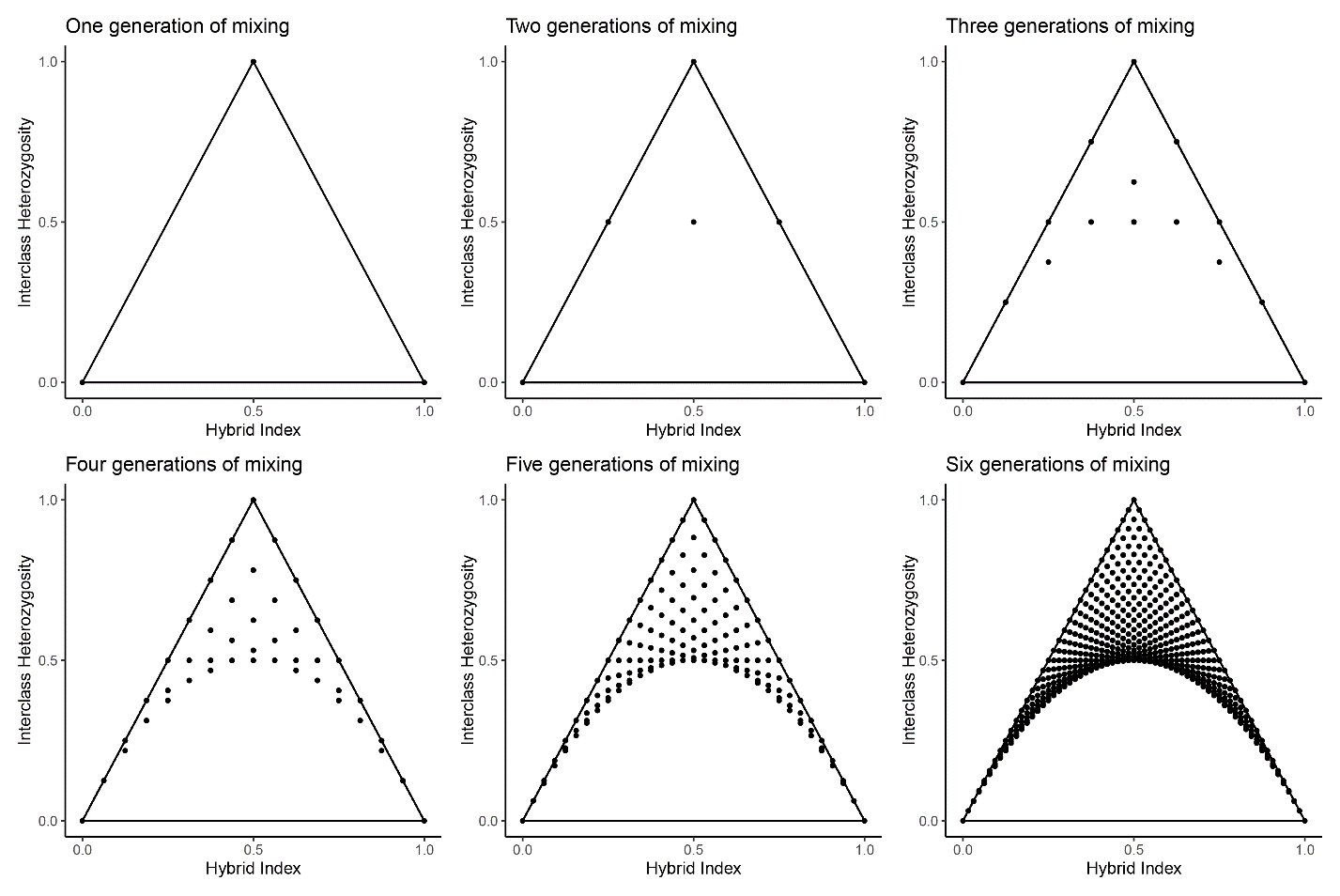

**Figure S1.** Calculation of possible coordinates of individuals on a triangle plot after one, and up to six, generations of all possible matings of all individuals in the previous generation, under Hardy-Weinberg Equilibrium.

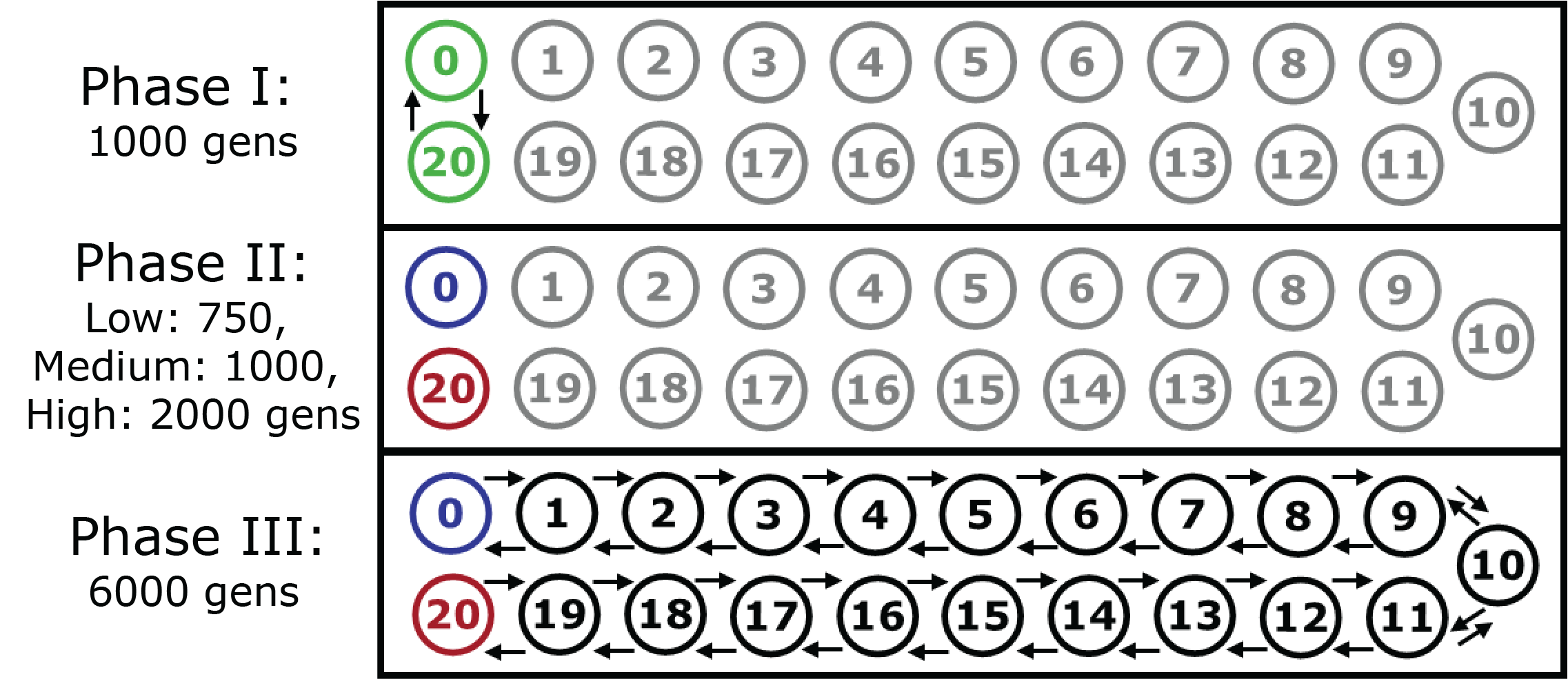

**Figure S2.** Schematic of the three phases of the simulations. In Phase I, individuals only occur in the green populations (p0 and p20), which are connected with a high migration rate, and no individuals exist in the gray populations. In Phase II, the parental populations (p0: blue; and p20: red) become allopatric, and evolve independently for 750, 1000, or 2000 generations. In Phase III, individuals expand from the parental populations until contact is made in the central population (pop 10). Migration between each population proceeds under a stepping-stone model.

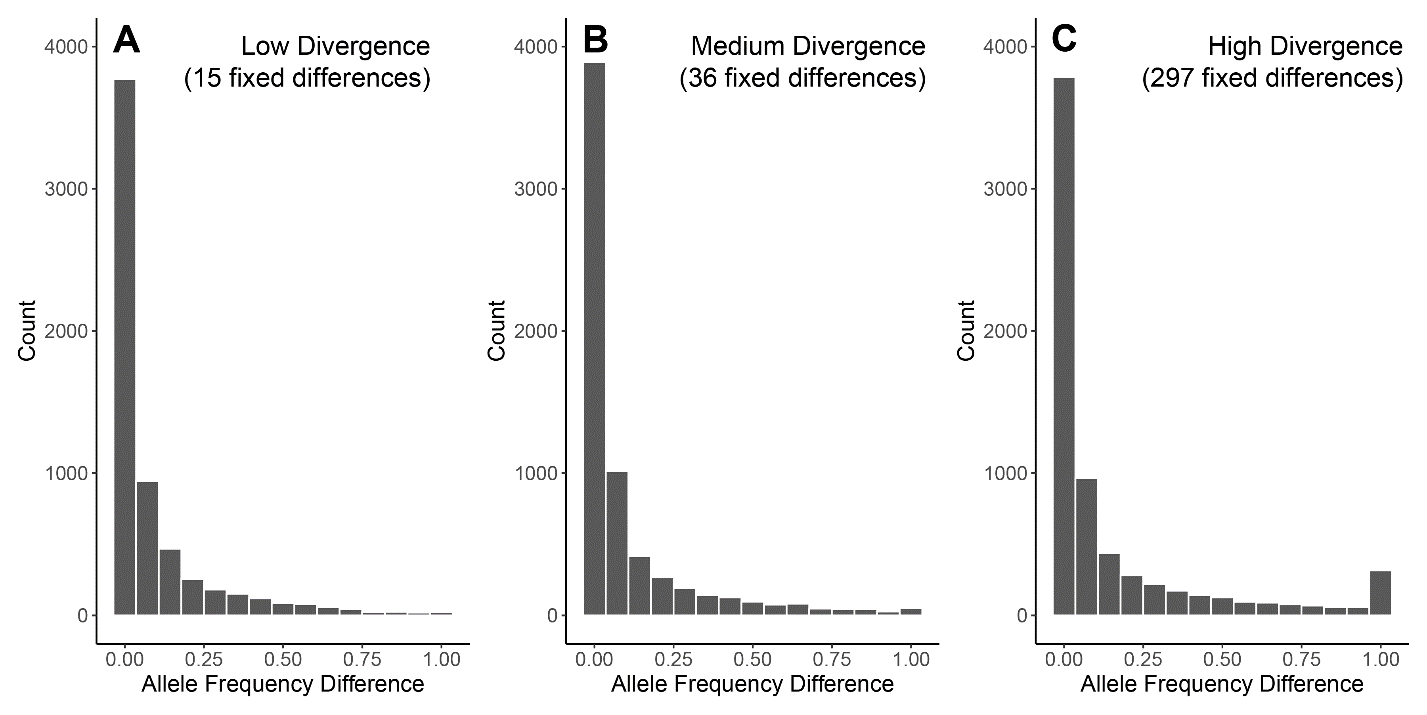

**Figure S3.** Spectrum of allele frequency differences between the parental populations in each simulation at the end of Phase II. There were 6251, 6511, and 6894 SNPs in the low **(A)**, medium **(B)**, and high **(C)** divergence simulations, respectively.

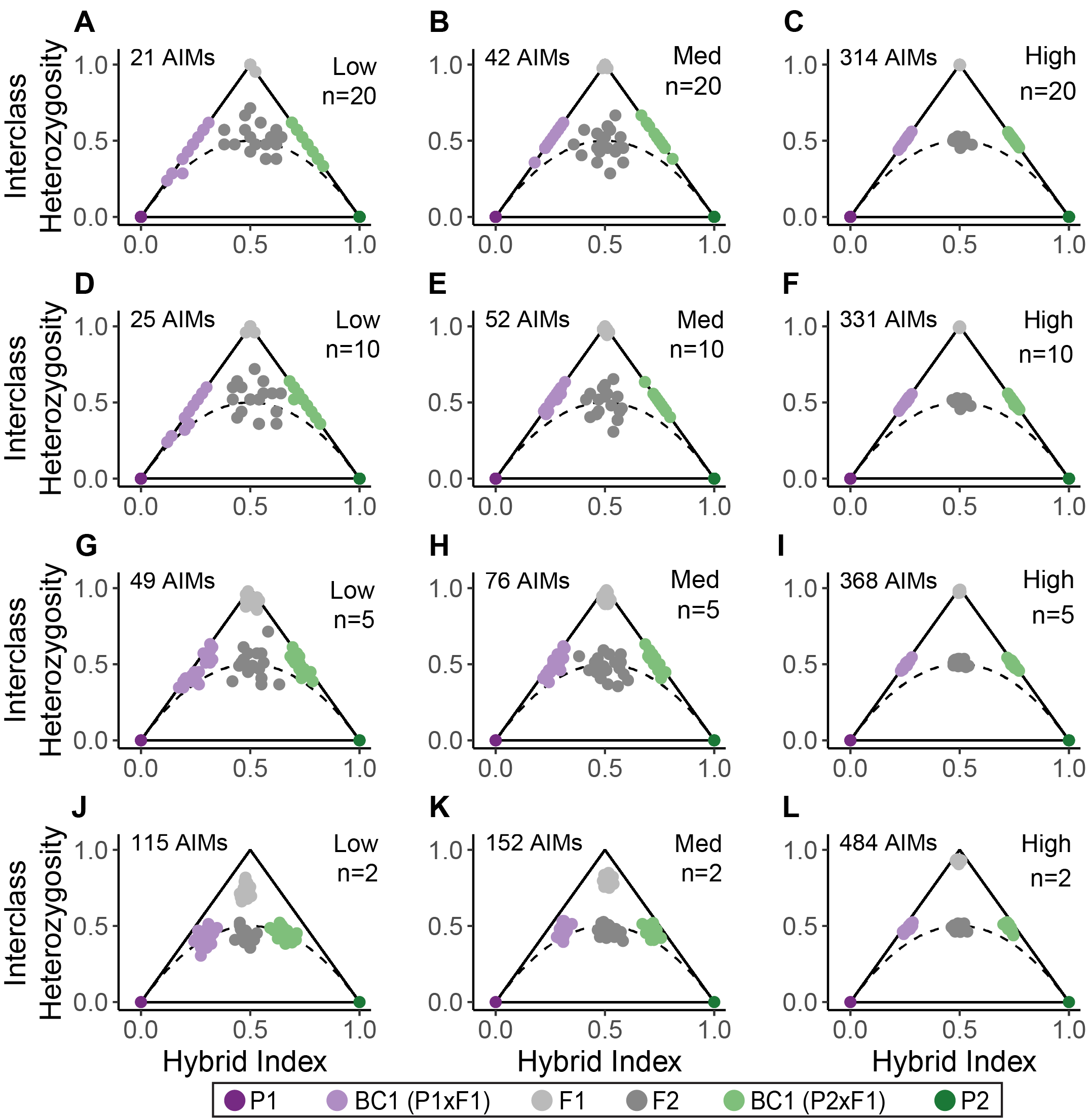

**Figure S4.** Triangle plots for known hybrids and parentals based on random samples of 20 **(A-C)**, 10 **(D-F)**, 5 **(G-I)**, or 2 **(J-L)** individuals (n) from each parental population. Sites that passed δ=1 in each sample were used as AIMs. The left column shows the simulation with low differentiation, the center column shows the simulation with medium differentiation, and the right column shows the simulation with high differentiation. Solid black lines indicate the possible space on a triangle plot, and the dotted black curve indicates the space below which individuals cannot occur, assuming Hardy-Weinberg Equilibrium.

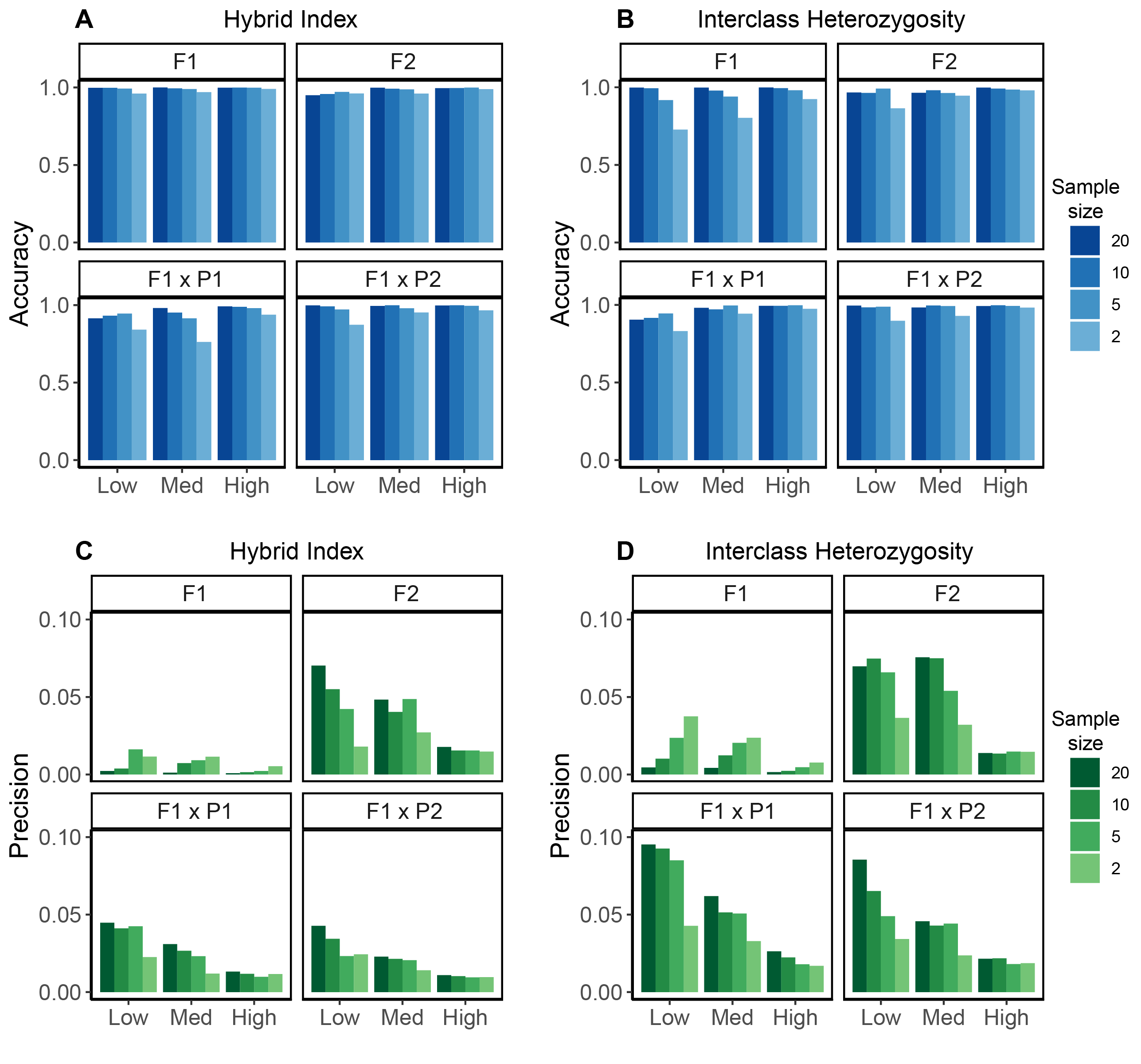

**Figure S5.** Accuracy **(A&B)** and precision **(C&D)** of hybrid index **(A&C)** and interclass heterozygosity **(B&D)** estimates based on random samples of 20, 10, 5, or 2 individuals from each parental population. Each simulation (low, medium, high) is shown on the x-axis. Accuracy and precision was measured for 20 individuals from each of the four hybrid classes (F1, F2, and the two first generation backcrosses) separately. Accuracy is reported as a percent, with 1 indicating 100% accuracy. Precision is reported as the average Euclidean distance of each observation within a class from the average of that class, such that smaller values indicate higher precision.

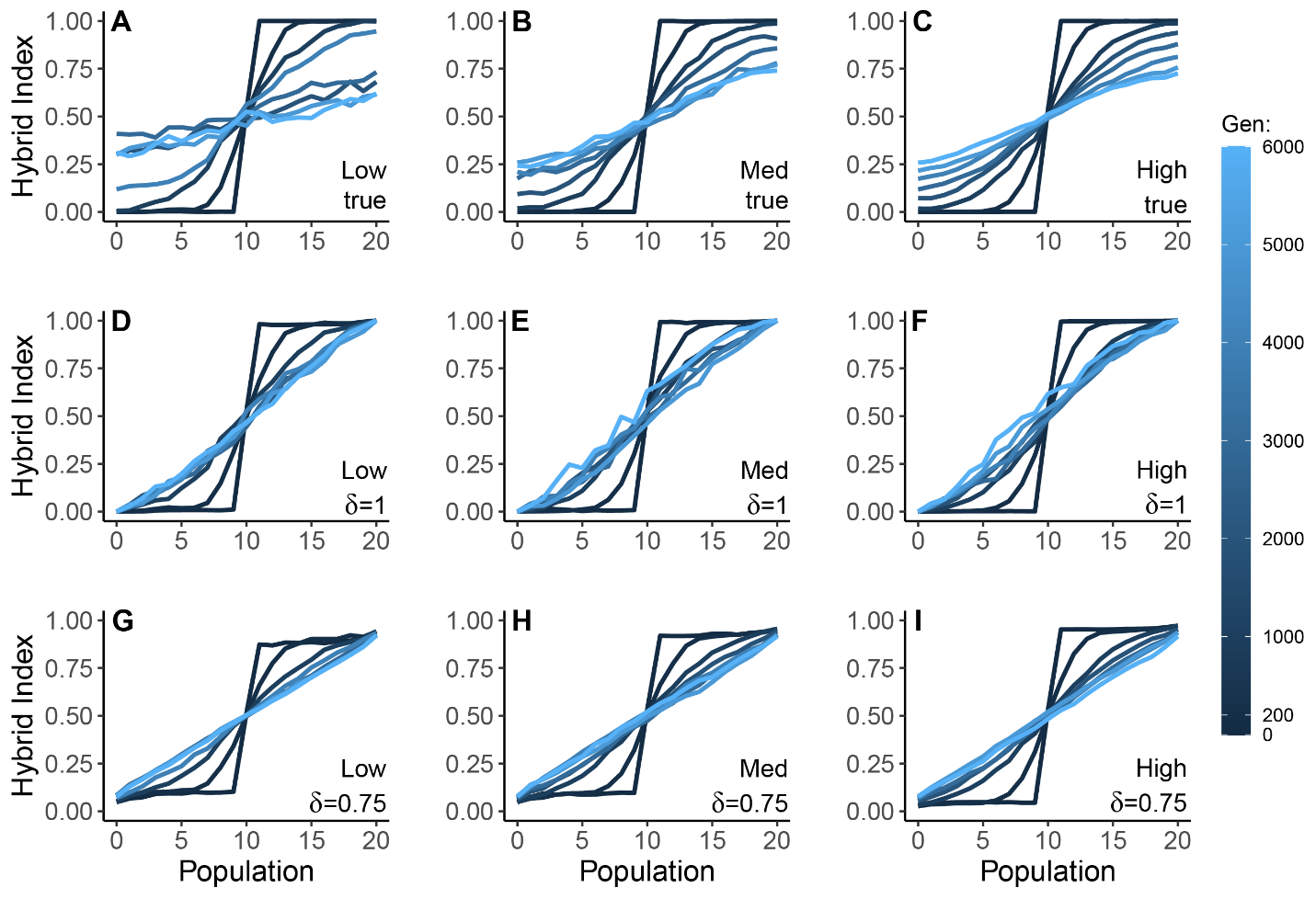

**Figure S6.** Hybrid index clines for true AIMs **(A-C)**, fixed differences identified within each sampled generation **(D-F)**, and AIMs that passed δ = 0.75 within each sampled generation **(G-I)**. The left column shows the simulation with low differentiation, the center column shows the simulation with medium differentiation, and the right column shows the simulation with high differentiation. Colors indicate the number of generations after initial contact, with generation 0 defined as the first generation in which all populations had at least 50 individuals during Phase III of the simulations.

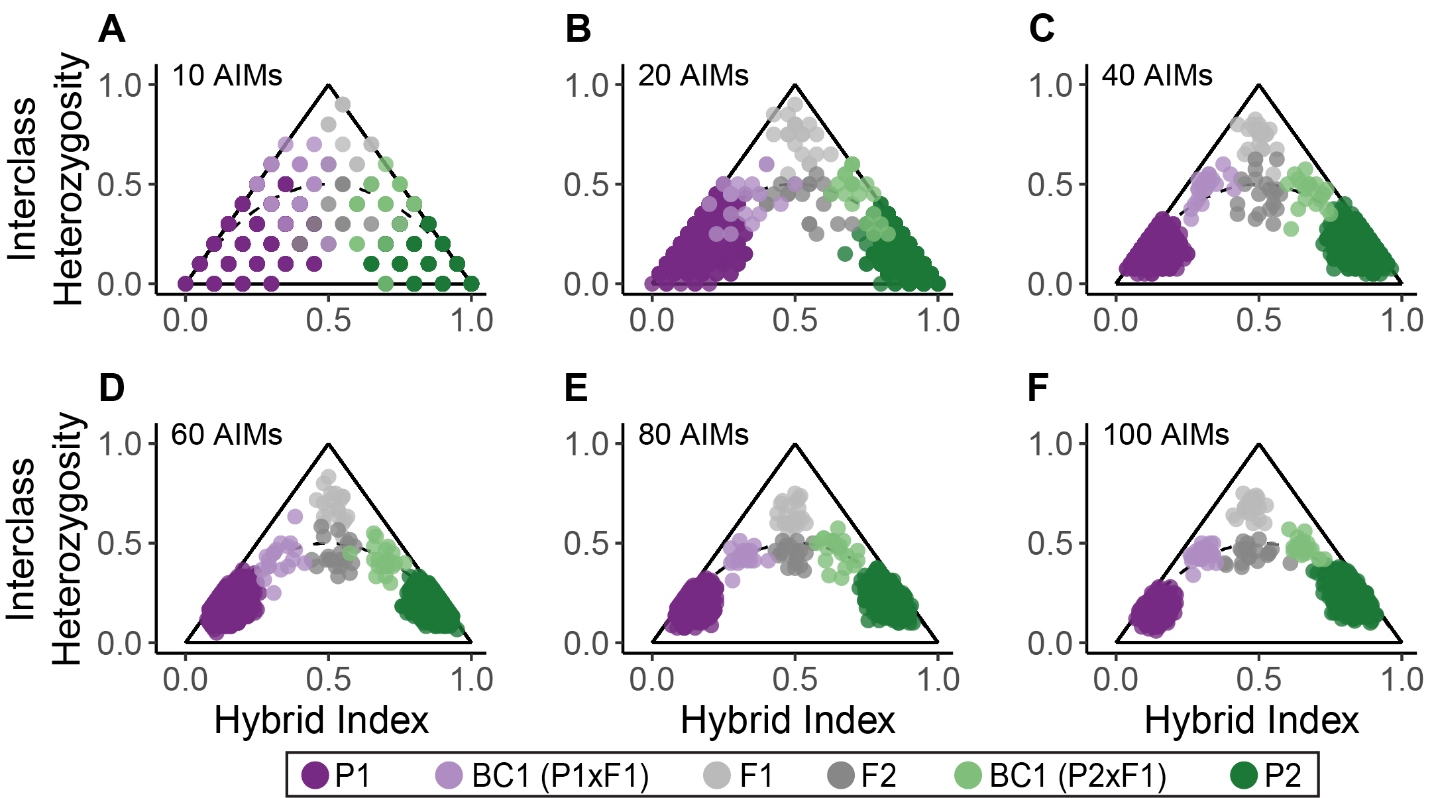

**Figure S7.** Triangle plots for known hybrids and parentals when divergence is low and few AIMs are identified. All plots are based on AIMs identified with the δ=0.5 threshold in the low divergence simulation. Of the AIMs identified with the δ=0.5 threshold (n=315), we randomly downsampled 10 **(A)**, 20 **(B)**, 40 **(C)**, 60, **(D)**, 80 **(E)**, and 100 **(F)** AIMs. Sampling AIMs in this way essentially models sequencing effort, and serves as a demonstration of the number of AIMs needed to accurately identify hybrid classes when parental divergence is low.

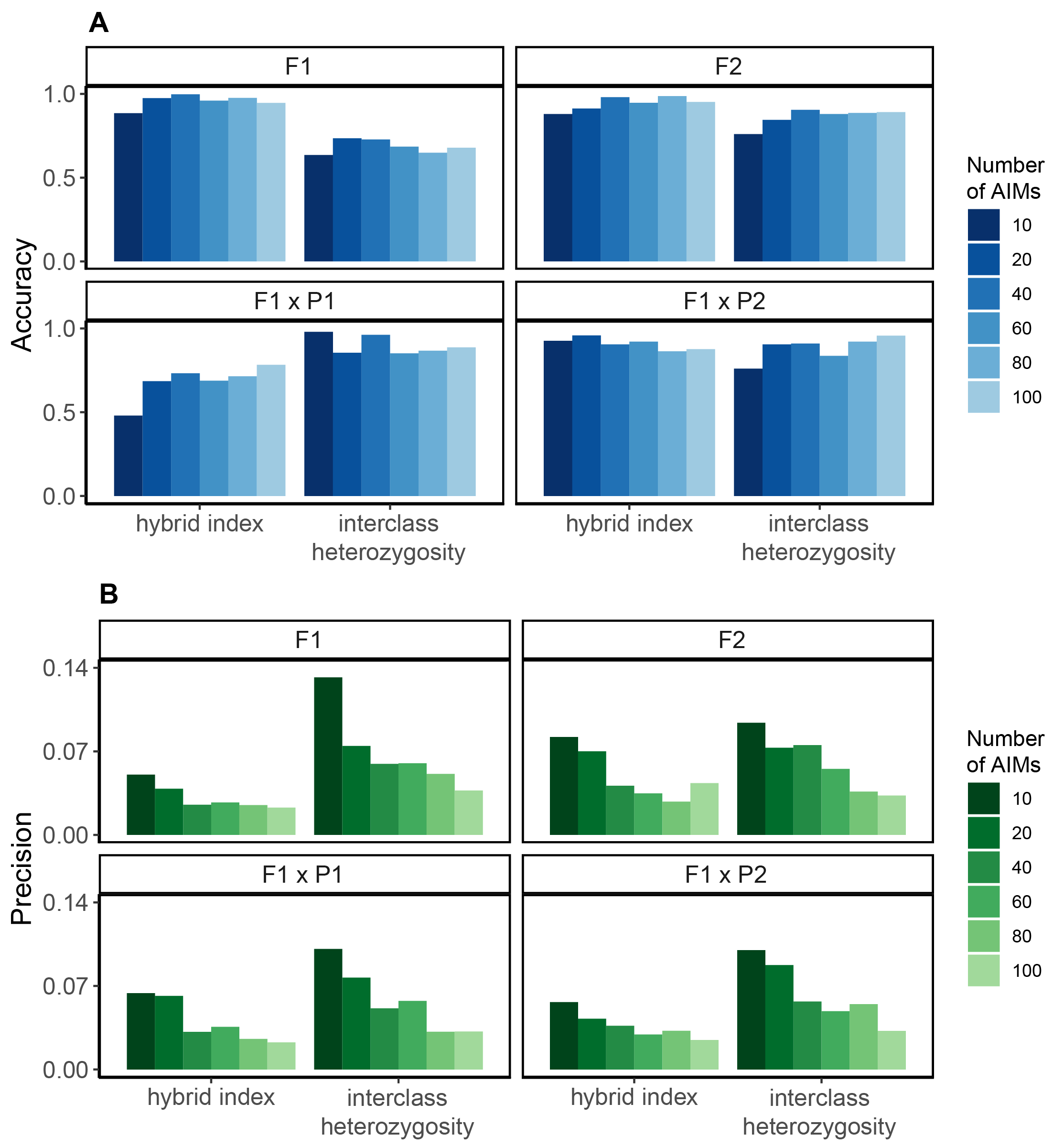

**Figure S8.** Accuracy **(A)** and precision **(B)** of hybrid index and interclass heterozygosity estimates when divergence is low and few AIMs are identified. AIMs were identified with the δ=0.5 threshold in the low divergence simulation, and were randomly downsampled to 10, 20, 40, 60, 80, and 100 AIMs. Accuracy and precision of the estimates was measured for 20 individuals from each of the four hybrid classes (F1, F2, and the two first generation backcrosses) separately. Accuracy is reported as a percent, with 1 indicating 100% accuracy. Precision is reported as the average Euclidean distance of each observation within a class from the average of that class, such that smaller values indicate higher precision.

**Table S1.** Descriptive statistics about the true allele frequencies of AIMs (δ=1) identified based on 200 random samples (n) of 20, 10, 5, and 2 parental individuals. “True δ<1” refers to the percent of AIMs that have δ=1 in the sample but δ<1 between the full populations. “True δ<0.95” refers to the percent of AIMs that have δ=1 in the sample but δ<0.95 between the full populations. Accuracy is defined as the average absolute difference between the expected value (true δ) and the observed value in the sample, divided by the expected value. We subtract that value from 1 to report the percent accuracy. “0.05 Quantile” indicates the value *x* for which 95% of identified AIMs have a true δ greater than *x*. Likewise “0.1 Quantile” and “0.25 Quantile” show this value at the 90% and 75% cutoffs, respectively.

| Sim | n | True δ<1 | True δ<0.95 | Accuracy | 0.05 Quantile | 0.10 Quantile | 0.25 Quantile |
| --- | --- | --- | --- | --- | --- | --- | --- |
| low | 20 | 28.8% | 3.9% | 99.3% | 0.96 | 0.98 | 0.99 |
| low | 10 | 45.1% | 17.7% | 97.8% | 0.89 | 0.92 | 0.98 |
| low | 5 | 63.8% | 42.1% | 93.1% | 0.72 | 0.78 | 0.89 |
| low | 2 | 85.7% | 76.1% | 76.4% | 0.39 | 0.47 | 0.63 |
| med | 20 | 19.1% | 2.9% | 99.5% | 0.97 | 0.98 | 1.00 |
| med | 10 | 33.6% | 14.3% | 98.2% | 0.90 | 0.94 | 0.98 |
| med | 5 | 53.1% | 35.9% | 94.3% | 0.77 | 0.82 | 0.91 |
| med | 2 | 77.1% | 67.5% | 81.4% | 0.44 | 0.54 | 0.69 |
| high | 20 | 4.6% | 0.7% | 99.9% | 1.00 | 1.00 | 1.00 |
| high | 10 | 9.7% | 3.8% | 99.5% | 0.96 | 1.00 | 1.00 |
| high | 5 | 18.8% | 12.0% | 98.2% | 0.87 | 0.94 | 1.00 |
| high | 2 | 38.0% | 31.9% | 91.9% | 0.58 | 0.70 | 0.89 |
